# PALP: An imaging method for detecting and quantifying polyunsaturated phospholipids via peroxidation

**DOI:** 10.1101/2020.04.11.037218

**Authors:** Yilong Zou, Emily T. Graham, Yuwei Huang, Wendy Salmon, Li Yu, Stuart L. Schreiber

**Author notes:** These authors contributed equally to this work (Yilong Zou, Emily T. Graham). Correspondence: Stuart L. Schreiber, Ph.D., Yilong Zou, Ph.D.

## Abstract

Polyunsaturated phospholipids are essential for multiple cellular functions; however, their uncontrolled peroxidation leads to ferroptosis. Here we describe *<u>p</u>*hotochemical *<u>a</u>*ctivation of membrane *<u>l</u>*ipid *<u>p</u>*eroxidation (PALP), which uses localized laser pulses to induce lipid peroxidation photochemically. While PALP bypasses enzymatic requirements for lipid peroxidation, the resulting BODIPY-C11-based signal is largely correlated with local polyunsaturated phospholipid concentration on membranes. This technique enables non-invasive reporting of lipid unsaturation levels and sensitivity to ferroptosis in live cells.

---

Ferroptosis, an iron-dependent, non-apoptotic cell death program, is a key contributor to tissue damage occurring in a variety of acute organ injury as well as chronic degenerative diseases^1^. Ferroptotic cell death is driven by aberrantly accumulated oxidative damage of membrane phospholipids possessing polyunsaturated fatty acyl (PUFA) chains, hence the cellular susceptibility to ferroptosis is highly dependent on cellular PUFA-phospholipid content and distribution. Recent research highlights that ferroptosis susceptibility varies dramatically among different lineages and cell states^2^. While targeting ferroptosis is emerging as an appealing strategy for overcoming certain diseases, the heterogeneity in ferroptosis susceptibility among different tissues and cell types will likely pose a challenge to achieve high efficacy with ferroptosis-inducing agents. Therefore, there is a pressing need for techniques that enable rapid and non-invasive assessment of cellular susceptibility to ferroptosis.

Multiple factors are known to contribute to the variable sensitivity to ferroptosis, including availability of reactive iron, PUFA-phospholipid concentrations in membranes, activities of lipid peroxidation-promoting enzymes, and cellular oxidative stress levels. Nonetheless, the rapid, irreversible and pervasive nature of ferroptosis has prevented detailed characterizations of the molecular processes of lipid peroxidation and ferroptotic cell death. Here, to enable visualization of lipid peroxidation at high spatio-temporal resolution, we use high-power laser pulses to induce local lipid peroxidation, the intensity of which is captured by fluorescent signals from oxidative modification of the BODIPY-C11 probe^3^. This technique, termed photochemical activation of membrane lipid peroxidation (PALP), bypasses the enzymatic reactions normally mediating lipid peroxidation in cells and provides an approximate estimation for the cellular PUFA-phospholipid levels.

To assess the dynamics of lipid peroxidation, we pre-treated the ferroptosis-susceptible 786-O clear-cell renal cell carcinoma (ccRCC) cell line^2^, with BODIPY-C11 (B-C11). B-C11 can be oxidized by membrane associated lipophilic reactive oxygen species and switches from orange (591 nm) to green (503nm) fluorescence after oxidation^4^. We applied 5 pulses of 405 nm light from a high-power laser source on a confocal microscope to a localized region in the cell over a 5 second period (**Fig. 1a**). The target cells were imaged for oxidized B-C11 (oxB-C11) signal for 25 seconds at 1 second intervals (**Fig. 1a**). We found that the 405 nm laser pulses induced strong oxB-C11 signal in 786-O cells immediately following the laser pulses, and the signal gradually decayed after reaching the peak intensity (normalized maximum intensity, PALP I-max) (**Fig. 1b-c, Supplementary Video 1**).

**Figure 1.**
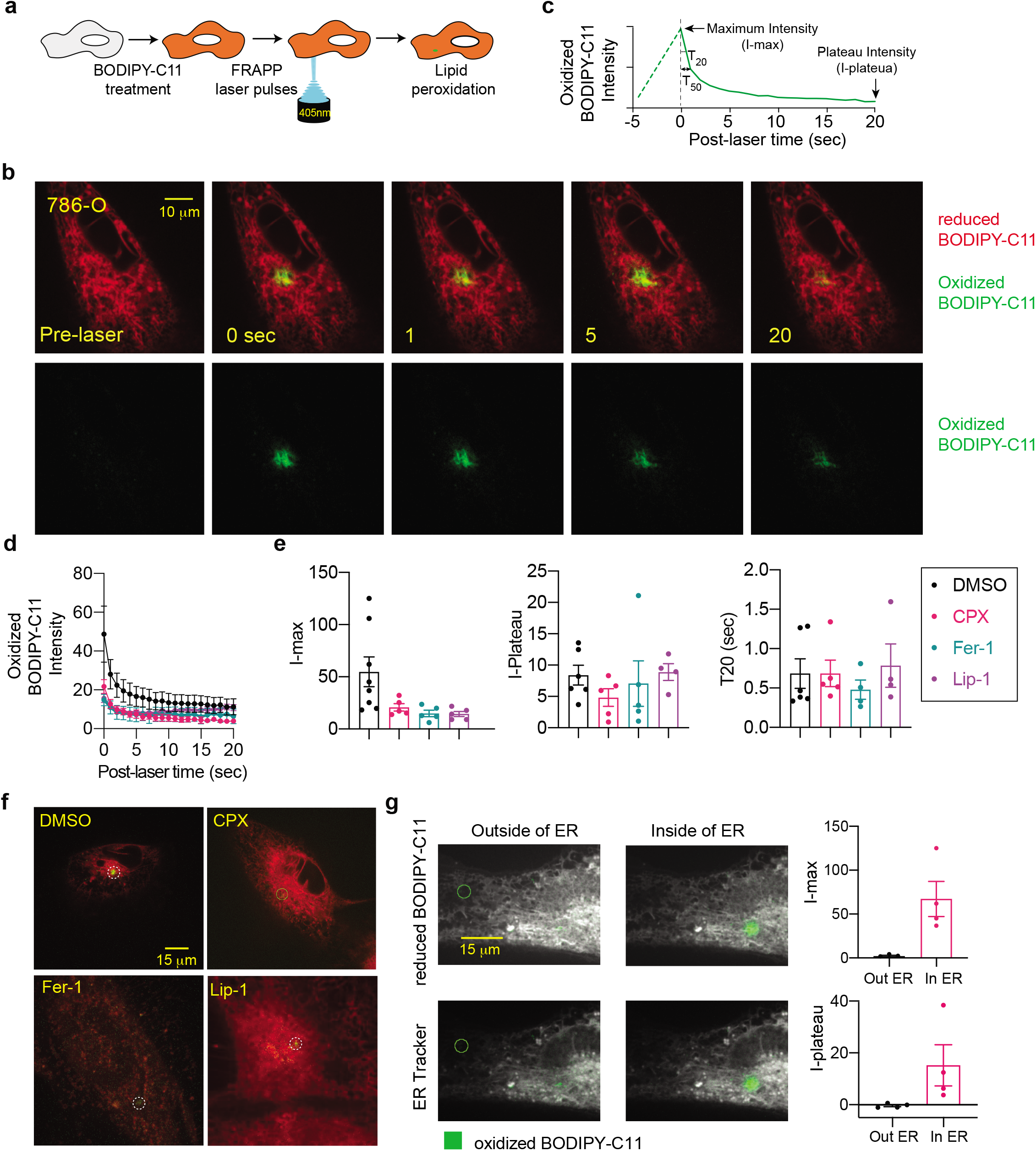
Targeted laser pulses induce localized lipid peroxidation in live cells. **a.** Schematic diagram describing the procedures of the photochemical activation of membrane lipid peroxidation (PALP) technique. BODIPY-C11 at a concentration of 5 μM was added to the cells approximately 30 min prior to imaging. Then a point on the cell was targeted by 5 pulses of the 405 nm laser. Each pulse lasts for 1 second. The 488 nm channel intensity was then recorded for 25 seconds at 1 second interval to monitor the oxidized BODIPY-C11 signal. **b.** Fluorescent images showing the time course of the reduced and oxidized BODIPY-C11 signal before and after the application of 405 nm laser pulses in wildtype 786-O cells. Scale bar indicates 10 μm. **c.** Schematic diagram showing the parameters used to describe the dynamics of lipid peroxidation signal in a representative PALP experiment. All oxBODIPY-C11 signals are normalized by deducing the pre-laser intensity from the detected signals. I-max, normalized maximal intensity, T20, time (sec) it takes to reach 20% reduction from I-max signal. T50, time (sec) it takes to reach 50% reduction from I-max signal. I-plateau, normalized signal intensity when the oxBODIPY-C11 signal largely plateaued. **d.** Quantifications of the PALP signal intensity from time-lapse imaging of 786-O cells treated with ciclopirox olamine (CPX), ferrostatin-1 (Fer-1) and liproxstatin-1 (Lip-1) and stimulated with laser pulses. 5 cells were measured for each condition, and error bars indicate mean±s.d. **e.** Box-scatter plots showing the PALP parameters of 786-O cells treated with ciclopirox olamine (CPX), ferrostatin-1 (Fer-1) and liproxstatin-1 (Lip-1). **f.** Representative fluorescent images showing the I-max (0 sec) post the application of laser pulses to 786-O cells treated with the indicated compounds. Scale bar indicates 15 μm. Green, oxidized BODIPY-C11 signal; red, reduced BODIPY-C11 signal. **g.** Image representation of endoplasmic reticulum (ER) staining and PALP lipid peroxidation intensities within and outside the ER in 786-O cells. Quantifications of I-max are on the right.

We next assessed whether the oxB-C11 signal is indeed resulted from lipid peroxidation. Key features of lipid peroxidation include its iron-dependence and ability to be quenched by lipophilic radical-trapping antioxidants (RTAs)^1,5^. Indeed, treatment with iron chelator ciclopirox olamine (CPX), or lipophilic RTAs and ferroptosis inhibitors ferrostatin-1 (Fer-1) and liproxstatin-1 (Lip-1)^5,6,7^ effectively reduced the PALP I-max signal in 786-O cells (**Fig. 1d-f, Supplementary Fig. 1a**), suggesting that PALP shares similar features of lipid peroxidation induced by chemical or genetic inhibition of glutathione peroxidase 4 (GPX4)^2,8^.

The subcellular origin of lipid peroxidation remains elusive, largely because the explosive signal expansion and prompt membrane damage after inducing lipid peroxidation at a whole cell level has prevented characterizations at high spatial-temporal resolution. Previous imaging analysis revealed that the initial lipid peroxidation signal co-localizes with markers of the endoplasmic reticulum (ER)^9^, highlighting the ER as the top candidate organelle where lipid peroxidation could initiate. To assess whether lipid peroxidation indeed occurs in the ER, we labeled the ER in 786-O cells with a fluorescent ER-tracker, and applied the laser pulses to regions both inside and outside of the ER network. We found that while laser pulses induced a strong signal when applied to ER-tracker marked areas, the signal is significantly weaker when applied to areas outside of ER (**Fig. 1g**). This result suggests that lipid peroxidation likely requires the ER for initiation. In our subsequent studies, we thus applied the laser pulses to the core ER structures, approximately reported by the presence of strong non-oxidized B-C11 signal.

Lipid peroxidation can spread non-enzymatically via autoxidation or enzymatically via the cytochrome P450 oxidoreductase (POR) and its associated redox partners in multiple neoplastic cell types including 786-O ccRCC cells^10^, and through arachidonate lipoxygenases *(ALOXs)* in certain other cellular contexts^11^. To determine whether laser-induced lipid peroxidation is dependent on POR activity, we applied laser pulses to *POR^-/-^* single-cell 786-O clones we previously generated^10^. As a control, we also analyzed cells that are depleted of KEAP1^12^, which promotes ferroptosis susceptibility in cancer cells by inducing degradation of NRF2, a master transcriptional regulator of cellular antioxidant enzymes^13,14^. As we found, POR-depletion did not significantly alter PALP I-max signal, which was modestly reduced in KEAP1-depleted cells (**Supplementary Fig. 1b**). This result suggests that POR activity is not required for PALP induction. This likely reflects that high-power laser is bypassing POR to initiate lipid peroxidation photochemically. On the other hand, this data also hints that POR is involved in early initiation, rather than propagation of cellular lipid peroxidation reactions. The modest effects of KEAP1-depletion suggest that activation of cellular antioxidant programs may only exert indirect effects in attenuating PALP. Together, these results indicate that PALP is largely a photochemical process that is independent of protein enzymatic activity. We hence speculated that the local PUFA-phospholipid concentration might be the major rate-limiting factor regulating the PALP I-max signal.

To investigate the relationship between cellular PUFA-phospholipid content and PALP signal, we first depleted acyl-CoA synthetase long chain family member 4 (ACSL4) in ccRCC cells using CRISPR/Cas9 (**Fig. 2a, Supplementary Fig. 2a**). ACSL4 catalyzes the conversion of long-chain fatty acids into fatty acyl-CoA, and is a key requirement for PUFA-phospholipid synthesis as well as a common mediator of ferroptosis susceptibility in multiple cellular contexts^2,15,16^. Previous lipidomic analysis showed that ACSL4-depletion selectively reduced the levels of cellular PUFA-phospholipid levels, with compensatory upregulation of PUFA-triacylglycerides (TAGs)^12^. As a result, ACSL4-depleted cells exhibited significantly lower susceptibility to GPX4 inhibition-induced ferroptosis (**Fig. 2b, Supplementary Fig. 2b**). Importantly, the PALP I-max signal was also significantly suppressed in ACSL4-depleted cells (**Fig. 2c-e**), suggesting that high levels of PUFA-phospholipids are necessary to potentiate strong PALP induction.

**Figure 2.**
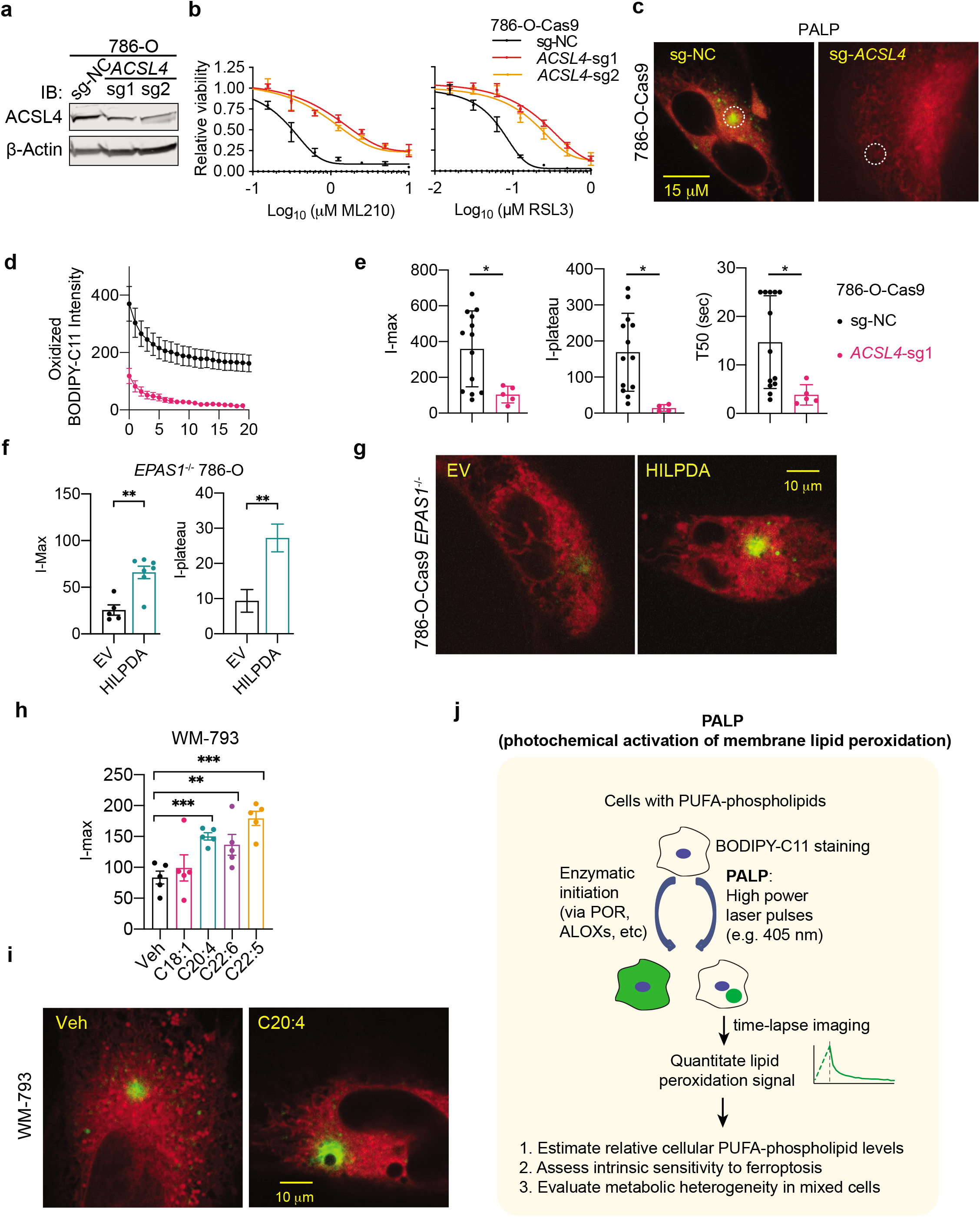
Photochemically activated lipid peroxidation signal correlates with polyunsaturated phospholipid levels. **a.** Immunoblot analysis of ACSL4 protein levels in 786-O cells expressing non-targeting negative control sgRNA (sg-NC) or *ACSL4*-targeting sgRNAs. β-actin was used as a loading control. **b.** Viability curves of 786-O cells expressing sg-NC or sgRNAs targeting *ACSL4* treated with indicated concentrations of GPX4 inhibitors ML210 or RSL3 for 48h. n=4; error bar, mean±s.d. **c.** Representative fluorescent images showing the I-max (0 sec) post the application of laser pulses to 786-O cells expressing sg-NC or *ACSL4*-targeting sgRNA. Green, oxidized BODIPY-C11 signal; red, reduced BODIPY-C11 signal. **d.** Quantifications of the PALP signal intensity from time-lapse imaging of 786-O cells expressing sg-NC or *ACSL4*-targeting sgRNA after laser activation. 5 cells were measured for each condition, and error bars indicate mean±s.d. **e.** Box-scatter plots showing the PALP parameters of 786-O cells expressing sg-NC or sgRNAs targeting *ACSL4.* Two-tailed unpaired T-test,*, p<0.05, **f.** Box-scatter plots showing the PALP parameters (I-max and I-plateau) of *EPAS1^-/-^* 786-O cells expressing empty vector (EV) or *HILPDA* cDNA. Two-tailed unpaired T-test,**, p<0.01. **g.** Representative fluorescent images showing the I-max (0 sec) post the application of laser pulses to *EPAS1^-/-^* 786-O cells expressing empty vector (EV) or *HILPDA* cDNA. Green, oxidized BODIPY-C11 signal; red, reduced BODIPY-C11 signal. **h.** Box-scatter plots showing the I-max of PALP signals in WM-793 melanoma cells treated with vehicle (veh) or indicated free fatty acids. Two-tailed unpaired T-test,**, p<0.01, ***, p<0.001. **i.** Fluorescent images showing the I-max (0 sec) post the application of laser pulses to WM-793 cells treated with vehicle (Veh, 5% BSA) or BSA conjugated arachidonic acid (C20:4). Scale bar indicates 10 μm. Green, oxidized BODIPY-C11 signal; red, reduced BODIPY-C11 signal. **j.** Schematic diagram summarising the applications of PALP for detecting and quantifying polyunsaturated phospholipid levels via photochemically-induced lipid peroxidation in live cells. PUFA, polyunsaturated fatty acyl-.

To explore whether PUFA-phospholipid upregulation is also sufficient to enable PALP induction, we used a previously established *EPAS1^-/-^* 786-O cell line model. In this model, ectopic overexpression of hypoxia-induced, lipid droplet-associated protein *(HILPDA),* a direct HIF-2α (encoded by *EPAS1*) target gene, selectively restores the cellular PUFA-lipidome, including both PUFA-phospholipids and PUFA-TAGs, in HIF-2α-depleted cells^2^. We found that HILPDA-overexpressing cells exhibited significantly stronger PALP I-max signal compared with empty vector-expressing cells (**Fig. 2f-g**), confirming that PUFA-lipids are rate-limiting for PALP induction.

We next used chemical approaches to complement the genetic modulation of PUFA-phospholipid levels. We pre-treated the human melanoma cell line WM-793^17^ with various synthetic fatty acids for three days prior to PALP application. These fatty acids include monounsaturated fatty acid (MUFA) oleic acid (OA, C18:1), and polyunsaturated fatty acids (PUFA) including arachidonic acid (AA, C20:4 ω6), docosapentaenoic acid (DPA, C22:5 ω3), and docosahexaenoic acid (DHA, C22:6 ω3). This experiment revealed that treatment with AA, DPA or DHA, but not OA significantly enhanced the PALP I-max signal; the extent of enhancement is correlated with the ferroptosis sensitization activities of these fatty acids^10^ (**Fig. 2h-i, Supplementary Fig. 2c**). In another cell line BFTC-909, a transitional renal cell carcinoma model, OA treatment even inhibited the PALP I-max signal (**Supplementary Fig. 2d-e**). The PALP-suppressive effect of MUFA is consistent with the recent identification of exogenous MUFAs as ferroptosis inhibitory agents^18^. Taken together, these results support positive correlations among cellular susceptibility to ferroptosis, the polyunsaturated-phospholipid levels, and the PALP I-max signal.

Recent research highlighted that PUFA-lipids play fundamental roles in shaping the ferroptosis sensitivity in various cellular contexts^2,9,19^. However, over-abundant PUFA-phospholipids in membranes create a vulnerability to ferroptosis induction. While single-cell lipidomics awaits to mature, there are limited tools to efficiently report the cellular PUFA-lipid levels in a cell. Here we showed that high-power laser-induced lipid peroxidation (PALP) shares similar chemical and biological properties as that induced by inhibition of GPX4 enzymatic activity. PALP bypasses the requirements for peroxidation-promoting enzymes including POR, and is largely dictated by the local PUFA-phospholipid levels in cellular membranes. Hence, the PALP technique provides a practical and efficient approach to non-invasively evaluate the PUFA-lipid abundances in live cells. We envision that PALP will be useful for studying the mechanisms of ferroptosis at high spatio-temporal resolution. As a proof-of principle, we used PALP to learn that lipid peroxidation requires the endoplasmic reticulum for initiation. Moreover, this technique could also be useful for rapidly evaluating the PUFA-lipid levels in a heterogeneous cell population and contribute to various fronts of lipid metabolism research.

## Methods

### Cell lines and culture conditions

786-O, 769-P and WM-793 cells were cultured with RPMI 1640 (Gibco) medium. BFTC-909 cells were cultured with DMEM (Gibco) medium. All cell lines were obtained from the Cancer Cell Line Encyclopedia (CCLE) distributed and authenticated by the Broad Institute Biological Samples Platform and Genetic Perturbation Platform. All culture media were supplemented with 10% fetal bovine serum (Gibco) and 1% penicillin/streptomycin. 786-O-Cas9 cells expressing *KEAP1-* or *ACSL4*-targeting sgRNAs were established and validated as previously described^12^. *POR^-/-^* single cell clones were generated and validated in prior studies^10^. HILPDA-expressing *EPAS1^-/-^* cells were generated as previously described^2^.

### Chemicals

ML210, RSL3, ferrostatin-1, liproxstatin-1 and ciclopirox olamine (CPX) were obtained from commercial sources (Sigma-Aldrich, SML0521, SML2234, SML0583, SML1414, and 1134030 respectively). CPX was used at 10 μM. Ferrostatin-1 was used at 5 μM. Liproxstatin-1 was used at 1 μM. BODIPY-C11 (Life Technologies) was reconstituted in DMSO, and added to the cells 30 min prior to live-cell imaging at 5 μM.

### Free fatty acid treatment

Oleic acid (OA, C18:1), arachidonic acid (AA, C20:4), docosapentaenoic acid (DPA, C22:5), and docosahexaenoic acid (DHA, C22:6) were purchased from Cayman Chemicals and conjugated with fatty-acid free BSA (Sigma-Aldrich) using previously described protocols^20^, and treated cells at 20 μM for 3 days prior to viability assays or imaging analysis.

### Viability assays

For cellular viability assays, cells were seeded in 384-well opaque white tissue culture and assay plates (Corning) at 1000 cells/well. 18-24 hours after seeding, cells were treated with compounds at the indicated concentrations for 48-72 hours. Cellular ATP levels were quantified using CellTiter-Glo Luminescence Assay (Promega) on an Envision multi-plate reader (PerkinElmer). Relative viability was normalized to the respective untreated condition of each cell line using RStudio and plotted in PRISM 8 (GraphPad software). For data presentation, the mean and standard deviation (s.d.) for the four biological replicates of each data point in a representative experiment is presented. Sigmoidal non-linear regression models were used to compute the regression fit curves.

### CRISPR/Cas9-mediated genome editing

Cells were engineered for constitutive Cas9 expression using the pLX-311-Cas9 vector (Addgene 96924), which contains the blasticidin S-resistance gene driven by the SV40 promoter and the SpCas9 gene driven by the EF1 α promoter. sgRNA sequences were cloned into the pLV709 doxycycline-inducible sgRNA expression vectors. Lentiviruses were generated from lentiviral Cas9 and sgRNA constructs in HEK-293T packaging cells. Lipofectamine 2000 (Life Technologies) was used as transfection reagents to deliver plasmids to cells following manufacturer’s instructions. Second generation packaging plasmids, including pMD2.G and pPAX2, was used for lentiviral production. Lentivirus titer was briefly assessed with Lenti-X Go-Stix Plus (TakaraBio). Target cells were infected with lentiviruses in the presence of 5 μg/mL of polybrene (Millipore). Depending on the vector, infected cells were selected with 2 μg/mL of puromycin or 10 μg/mL of Blasticidin S and propagated for further analysis. Cells transduced with doxycycline-inducible constructs were treated with 1 μg/ml of doxycycline (Sigma-Aldrich) for 7-14 days prior to gene-knockout validation using immunoblotting. Sequences of sgRNAs used in CRISPR experiments are: *ACSL4-sg1,* GCATCATCACTCCCTTAGGT; *ACSL4-sg2,* GTGTGTCTGAGGAGATAGCG.

### Immunoblotting

Adherent cells were briefly washed twice with ice-cold PBS and lysed with 1 % SDS lysis buffer containing 10 mM EDTA and 50 mM Tris-HCl, pH 8.0. Lysates were collected, briefly sonicated, then incubated at 95 °C for 10 min and the protein concentrations were determined using the BCA Protein Assay kit (Pierce) following manufacturer’s instructions. Calibrated samples were diluted with 4x lithium dodecyl sulfate (LDS) sample buffer (Novus), separated by SDS–PAGE using NuPAGE 4–12% Bis-Tris protein gels (Novus), and transferred to nitrocellulose or PVDF membranes by an iBlot2 protein-transfer system (Thermo Fisher Scientific). Membranes were blocked with 50% Odyssey blocking buffer (LiCor) diluted with 0.1% Tween-20-containing Tris buffered saline (TBST) and immunoblotted with antibodies against ACSL4 (Abcam, ab155282, produced in rabbit, used at 1:1000 dilution) and β-Actin (8H10D10, no. 3700 and 13E5, no. 4970, Cell Signaling Technologies, used at 1:5,000 dilution). Membranes were then washed with TBST and incubated with IRDye 800CW goat-anti-Rabbit or 680RD donkey-anti-Mouse secondary antibodies (LiCor). Immunoblotting images were acquired on an Odyssey equipment (LiCor) according to the manufacturer’s instructions, and analyzed in the ImageStudio software (LiCor). β-Actin was used as a loading control.

### Laser imaging and data analysis

Imaging was performed on an Andor Revolution Spinning Disk Confocal, FRAPPA and TIRF microscope. Prior to high-power laser application, steady state images were acquired to visualize the distributions of reduced and oxidized BODIPY-C11 in cells respectively. Each confocal laser was set at 3mHz and standard gain with a 200 ms exposure. FRAPPA bleaching was done with a 405 nm laser for 5 pulses and the 488 nm confocal channel was used to collect images for another 25 seconds following the laser pulse. Images are acquired using the Metamorph software associated with the equipment. Subsequently, image analysis was performed using Fiji ImageJ (1.52P). ROIs were imported from metamorph to locate laser bleaching regions. Each series measurement was analyzed using ImageJ ROI manager multi-measurement tool at a 10 px radius around the region of interest. Data was normalized to time point before bleaching occurred and the resulting plot was fit with a non-linear regression to find I-max (Y0), T50, Plateau and T20 in Prism 8 (GraphPad).

### Software and statistical analysis

Data are generally expressed as mean±*s.d.* unless otherwise indicated. No statistical methods were used to predetermine sample sizes. Statistical significance was determined using a two-tailed, unpaired student’s T-test using Prism 8 software (GraphPad Software) unless otherwise indicated. Statistical significance was set at p < =0.05 unless otherwise indicated. Figures are finalized in Adobe Indesign and Illustrator.

## Data and Code Availability Statement

Raw videos showing the dynamics of lipid peroxidation signal in wildytpe 786-O cells are provided as a **Supplementary Video**. All original data and computational code that support the findings of this study are available upon request.

## Acknowledgements

This work was supported in part by the NCI’s Cancer Target Discovery and Development (CTD^2^) Network (grant number U01CA217848, awarded to S.L.S.), and in part by the National Institute of General Medical Sciences (grant number R35GM127045, awarded to S.L.S.). Y. Zou was supported by the National Cancer Institute of the National Institutes of Health under Award Number K99CA248610.

## Author contributions

Y.Z. initiated and conceived the project with input from Y.H. and L.Y. S.L.S. supervised the project. E.G. and Y.Z. performed the experiments and data analyses. W.S. assisted the imaging analyses.

## Additional information

Further information and requests for resources and reagents should be directed to the corresponding authors Stuart L. Schreiber or Yilong Zou.

## Supplementary Figure Legends

**Supplementary Figure 1.**
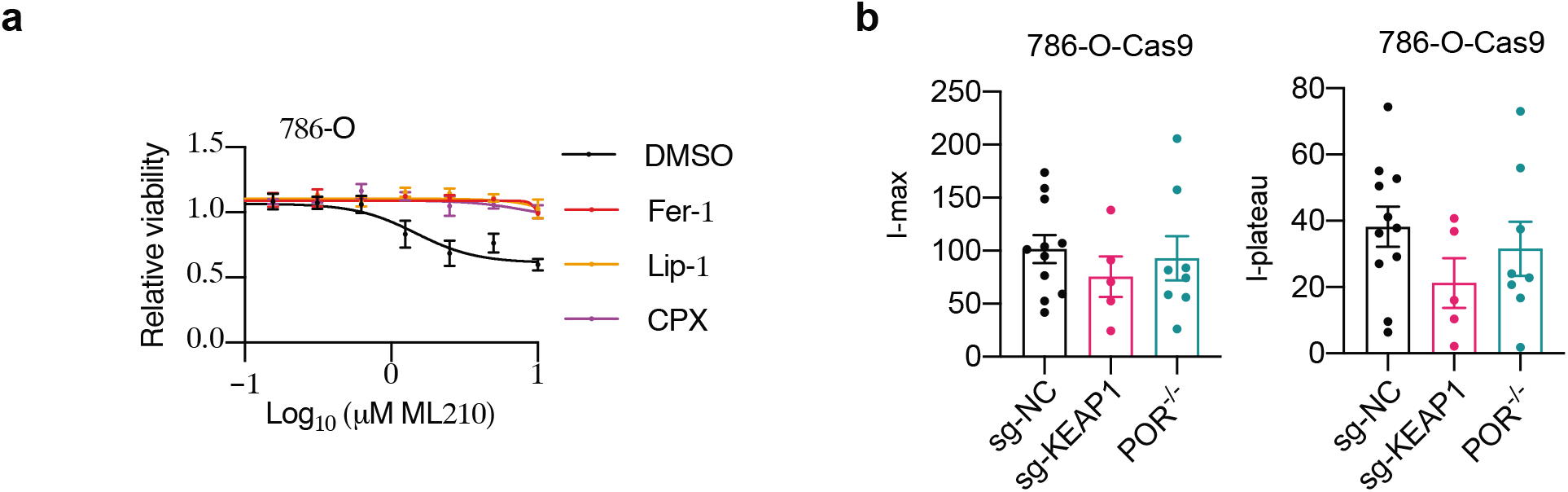
Targeted laser pulses induce localized lipid peroxidation in live cells. **a.** Viability curves for 786-O cells treated with ciclopirox olamine (CPX), Lip-1, or Fer-1 and ML210 for 48h. n=4; error bar, mean±s.d. **b.** Box-scatter plots showing the PALP parameters (I-max and I-plateau) of 786-O-Cas9 cells expressing sg-NC or *sg-KEAP1,* or a *POR^-/-^* 786-O single-cell clone.

**Supplementary Figure 2.**
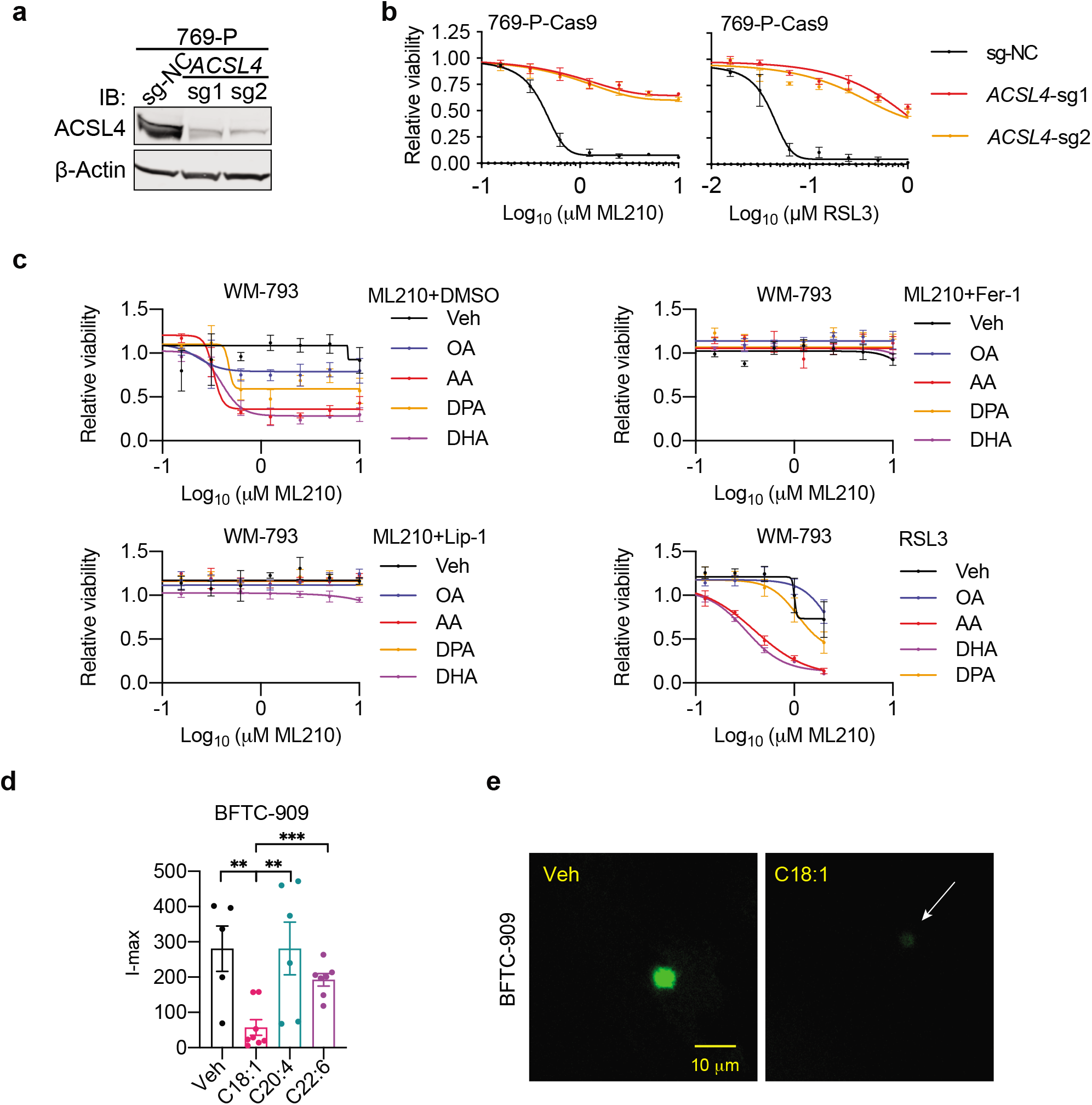
PALP signal correlates with cellular polyunsaturated phospholipid levels. **a.** Immunoblot analysis of ACSL4 protein levels in 769-P cells expressing negative control sgRNA (sg-NC) or *ACSL4*-targeting sgRNAs. β-actin was used as a loading control. **b.** Viability curves of 769-P cells expressing sg-NC or *ACSL4*-targeting sgRNAs treated with indicated concentrations of GPX4 inhibitors ML210 or RSL3 for 48h. n=4, error bar, mean±s.d. **c.** Viability curves for WM-793 cells treated with vehicle or indicated fatty acids, together with DMSO, Lip-1 or Fer-1 and indicated concentrations of ML210 or RSL3 for 48h. n=4, error bar, mean±s.d. **d.** Box-scatter plots showing the I-max of PALP signals in BFTC-909 transitional renal cell carcinoma cells treated with indicated free fatty acids. Two-tailed unpaired T-test,**, p<0.01, ***, p<0.001. **e.** Representative fluorescent images showing the I-max (0 sec) post the application of laser pulses to BFTC-909 cells treated with vehicle or oleic acid (C18:1). Green: oxidized BODIPY-C11 signal.

**Supplementary Video 1.** TIme-lapse imaging of oxidized BODIPY-C11 signal in 786-O cells stimulated with PALP. White signal, oxidized BODIPY-C11 signal. The time scale is accelerated at 3x speed.

